# Functional divergences of natural variations of *TaNAM-A1* controlling leaf senescence initiation during wheat grain filling

**DOI:** 10.1101/2023.04.22.537891

**Authors:** Longxi Zhou, Chuncai Shen, Wan Teng, Xue He, Xueqiang Zhao, Yanfu Jing, Zhixiong Huang, Yiping Tong

## Abstract

Leaf senescence is an essential physiological process related to grain yield potential and nutritional quality. Green leaf duration (GLD) after anthesis directly reflects the leaf senescence process and exhibits large genotypic differences in common wheat; however, the underlying gene regulatory mechanism is still lacking up to now. Here, we report *TaNAM-A1* as the causal gene of major loci *qGLD-6A* for GLD during grain filling by map-based cloning. The role of TaNAM-A1 in regulating leaf senescence, spike length, and grain size was proved by transgenic assay and TILLING mutants analyses. Furthermore, the functional divergences among *TaNAM-A1* three haplotypes were systematically evaluated. Wheat varieties with *TaNAM-A1d* (containing two mutations in CDS of *TaNAM-A1*) had longer GLD and advantages in yield-related traits than those with the wild type *TaNAM-A1a*. All three haplotypes were functional in transactivating the expression of genes involved in macromolecular degradation and mineral nutrient remobilization, with TaNAM-A1a the strongest activity and TaNAM-A1d the weakest. TaNAM-A1 modulates the expression of *TaNAC016-3A* and *TaNAC-S-7A* to trigger senescence initiation. TaNAC016-3A enhances TaNAM-A1 transcriptional activation ability by protein-protein interaction. Our study provides new insights into fine-tuning the leaf functional period and grain yield formation for wheat breeding under different geographical climatic conditions.

## Introduction

For wheat as an annual crop, monocarpic senescence is the final stage of its development and determines grain yield potential and nutritional quality. Monocarpic senescence is a complex process that refers to the programmed cell death with cellular macromolecular degradation in vegetative organs, and nutrients and amino acids transportation to developing grain (Ricachenevsky et al., 2013). Loss of chlorophyll is the visible reflection of leaf senescence (Thomas & Ougham, 2014). Recording the greenness or chlorophyll content changes of leaves is a typical way to monitor the senescence process over time (Chapman et al., 2021a). The “stay-green” phenotype, exhibiting long greenness duration of leaves, stems, and spikes, is often selected by breeders for its association with prolonged photosynthetic activity and a yield advantage, especially under stress environments such as drought, heat, and nutrients deficiency conditions (Richards, 2000; Bogard et al., 2011; Derkx et al., 2012; Naruoka et al., 2012). But not all the stay-green phenotypes are of the same superiority in yield and quality (Gous et al., 2013; Gregersen et al., 2013; Borrill et al., 2015). Rapid and efficient senescence is of equal importance, which ensures sufficient nutrient remobilization in a shorter growing season and is associated with higher grain protein and micronutrient concentration (Gong et al., 2005; Gregersen et al., 2008; Waters et al., 2009). One strategy to maximize individual grain yield and nutritional quality is to delay the senescence onset but speed the senescence rate (Wu et al., 2012; Xie et al., 2016). However, senescence is a tightly regulated process, with coordinated modulation of multiple pathways referring to the expression or component changes of thousands of genes (Lee & Masclaux-Daubresse, 2021). Therefore, uunderstanding the molecular mechanisms of senescence regulation is important for us to breed improved grain productivity and quality by balancing the senescence process with suitable grain filling duration in a certain environment (Distelfeld et al., 2014).

Many studies have revealed the senescence-associated gene transcriptional networks during leaf aging in many species such as Arabidopsis, rice, maize, and wheat (Gregersen & Holm, 2007; Breeze et al., 2011; Sekhon et al., 2012; Leng et al., 2017; Borrill et al., 2019). The transcription factors (TFs) play crucial roles in regulating gene expression in the senescence process. To date, several TFs have been identified as the regulator of leaf senescence in wheat. TaWRKY7 (Zhang et al., 2016), TaWRKT42-B (Zhao et al., 2020), and TaWRKY40-D (Zhao et al., 2020) have been shown to induce the expression of senescence-associated genes (SAGs) and accelerate senescence when overexpressed in leaves. NAC TFs also have critical function in controlling senescence progression. For example, TaSNAC11-4B functions as a positive regulator of senescence (Zhang et al., 2020), whereas TaNAC-S functions as a negative regulator (Zhao et al., 2015). The most well-studied NAC TF gene in wheat is *NAM-1/Gpc-1*. The B genome homolog in tetraploid wheat (*Triticum turgidum ssp. dicoccoides*), *TtNAM-B1*, known as *GPC-B1* was first cloned in identifying a major quantitative trait locus (QTL) for grain protein, zinc and iron content and accelerates senescence in flag leaves (Uauy et al., 2006). However, in most hexaploid common wheat varieties, *TaNAM-B1* is a non-functional copy or completely deleted. The functional homoeologous copies are *TaNAM-A1* and *TaNAM-D1* (Uauy et al., 2006), located on chromosomes 6A and 6D, respectively. The functions of *TaNAM-A1* and *TaNAM-D1* in regulating leaf senescence and grain mineral contents have previously been characterized by using EMS and TILLING mutation and transgenic modification of *TaNAM-A1* and *TaNAM-D1* expression. Silencing all the *NAM-1* homologs by RNA interference (*TaNAM*-RNAi) in hexaploid wheat significantly delayed senescence by prolonging the maintenance of chloroplast structure and high enzymatic antioxidant activity (Checovich et al., 2016), but reduced grain protein and micronutrients content (Uauy et al., 2006; Waters et al., 2009; Cantu et al., 2011; Guttieri et al., 2013). A similar phenotype was also presented in *gpc-1* TILLING mutants (Avni et al., 2014; Pearce et al., 2014). Subsequently, intensive studies concentrated on the function of A genome homolog *TaNAM-A1* (Pearce et al., 2014; Harrington et al., 2019; Chapman et al., 2021b) and its natural variations among modern wheat varieties (Cormier et al., 2015; Alhabbar et al., 2018). It has been revealed that the loss-of-function mutant of *TaNAM-A1* itself is sufficient to cause a significant delay in senescence of leaves (Avni et al., 2014; Pearce et al., 2014), and peduncles (Harrington et al., 2019). Several more recent studies focusing on transcriptome analyses of *TaNAM*-RNAi lines and TILLING mutants have shed light on the genetic mechanism of *NAM-1* controlling leaf senescence and nutrient remobilization (Cantu et al., 2011; Pearce et al., 2014; Harrington et al., 2020). However, the contribution of natural variation at *TaNAM-A1* locus to leaf senescence has yet to be illustrated genetically. Furthermore, the directly downstream regulation pathway and the functional divergence of natural variations of *TaNAM-A1* are still lacking up to now.

In this study, we identified *TaNAM-A1* as the causal gene for a major QTL *qGLD-6A* conferring stay-green trait during the grain filling stage by map-based cloning. TaNAM-A1 controls the leaf senescence initiation by directly up-regulating the genes involved in biomacromolecule degradation and iron and zinc remobilization, and is also involved in the regulation of spike length and grain size. The natural variations of *TaNAM-A1* lead to functional divergence in transactivating downstream genes. These findings will provide potential gene resources for fine-tuning grain filling duration purposefully by breeders.

## Results

### Map-based cloning of *qGLD-6A*

Previously, we developed a pair of QTL isolines, 178A and 178B, segregating for *qTaLRO-B1* alleles by using the heterogeneous inbred family analysis method (Cao et al., 2014). 178A and 178B showed differences in leaf senescence dynamics and yield-related traits under field conditions. During grain-filling, leaf relative chlorophyll value of each line was measured by SPAD-502 meter (Konica-Minolta, Japan) at different DAA. The duration from flowering date to the day when 50% flag leaves SPAD value lower than 35 was defined as green leaf duration (GLD) based on previous studies (Vijayalakshmi et al., 2010; Naruoka et al., 2012).178A exhibited longer green leaf duration (GLD) and delayed onset of leaf senescence compared to 178B under both low-nitrogen (LN) and high-nitrogen (HN) conditions (Figure S1a-c).

To understand the underlying genetic regulation of leaf senescence in wheat, we conducted QTL mapping for GLD at grain filling stage using RILs population derived from the cross between 178A and 178B and genotyping by the 90K Infinium iSelect SNP array (Figure S1d; Table S1). Consequently, three major QTLs for GLD on chromosomes 1A, 1B and 6A were detected under both LN and HN conditions in two growing seasons (Figure S1e; Table S2). The loci *qGLD-6A* with the highest LOD score can explain more than 50% phenotypic variation, and correspond to the region flanked by two SNP markers IWB6447 and IWB11771 (Figure 1a). Then we screened recombinants and residual heterozygous lines within this region (Figure 1b and 1c) and finally fine-mapped *qGLD-6A* to a 1-Mb region. Seven genes with high confidence were annotated within this region (Figure 1d), among which *TraesCS6A02G108100* and *TraesCS6A02G108300* are specifically expressed in leaves and were supposed to be the candidate genes (Table S3). We cloned *TraesCS6A02G108100* and *TraesCS6A02G108300* genomic sequences in 178A and 178B. There is one synonymous variation in the coding sequence of *TraesCS6A02G108100* between 178A and 178B (Figure S2). While in the open reading frame of *TraesCS6A02G108300*, 178A contained a 731C>T SNP leading to a missense variant (p.Ala171Val) and a deletion at 1518A leading to a frameshift variant (p.Gln396fs), compared to 178B (Figure 1e). *TraesCS6A02G108300* encodes a NAC transcription factor, which was previously designed as *TaNAM-A1*/*Gpc-A1*. *TaNAM-A1* is a homolog to *TaNAM-B1/Gpc-B1* (Uauy et al., 2006), controlling leaf senescence and grain protein concentration. Therefore, we determined *TaNAM-A1* as the candidate gene for *qGLD-6A.* Moreover, no significant difference in *TaNAM-A1* expression level was detected between 178A and 178B (Figure 1f). In a previous study, four haplotypes of *TaNAM-A1* were characterized based on the SNP and InDel in the coding sequence in hexaploid bread wheat (Cormier et al., 2015). The 178B allele belongs to the wild type *TaNAM-A1a*, and 178A allele belongs to the *TaNAM-A1d* haplotype.

**Figure 1.**
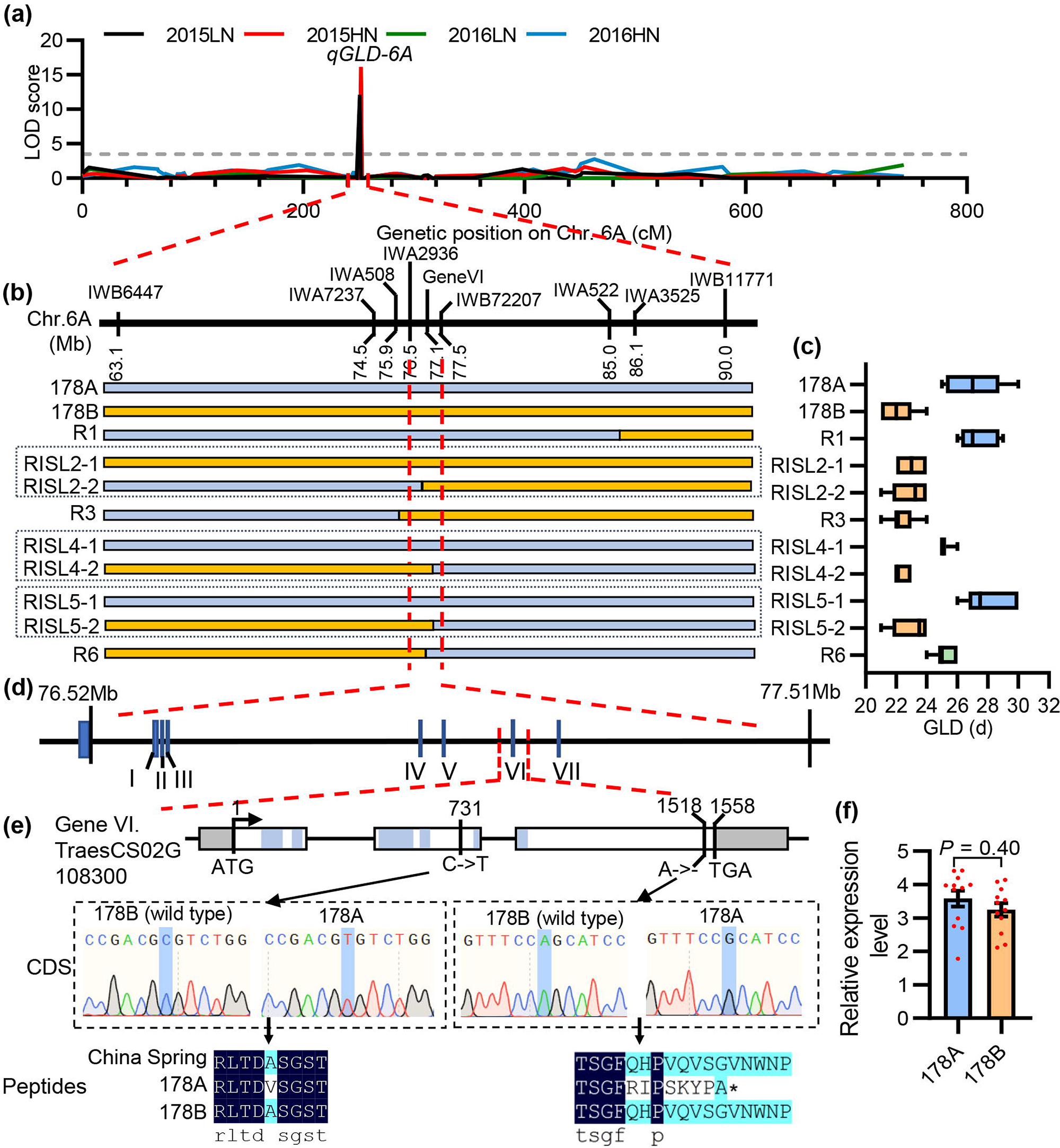
Map-based cloning of *qGLD-6A*. **(a)** Fine-mapping of *qGLD-6A*. Grey dashed line represents the LOD threshold was set to 3.5. **(b)** Graphical genotypes of critical recombinants used for fine-mapping of *qGLD-6A*. Blue and orange bars represent homozygous for 178A and 178B alleles respectively. Dashed line boxes indicate these two recombinant inbred sister lines (RISL) were derived from the same heterozygous recombinant inbred lines. **(c)** Green leaf duration of 178A,178B and their hybrids showed in **(b)** under low nitrogen conditions. Lines in the box plots indicate the median. More than six biological replicates for each line. **(d)** Seven genes with high confidence were annotated within 1 Mb region flanked by closest recombination events. **(e)** Sequence variations of *TraesCS6A02G108300* in 178A and 178B compared with reference genome China Spring. Numbers above represent physical locations within gene genomic sequence. The 5 conserved NAC subdomains were highlighted in blue. **(f)** qRT-PCR analysis of *TaNAM-A1* expression level in flag leaves of 178A and 178B at 17 DAA. Data were represented as mean ± SE, *n* = 12. Means were compared between 178A and 178B by independent sample t-test.

### TaNAM-A1 functions as a positive regulator of leaf senescence and is involved in spike elongation and grain size regulation

To validate the role of *TaNAM-A1* involved in leaf senescence regulation, we firstly constructed *TaNAM-A1a* overexpression lines in wheat variety KN199 (*TaNAM-A1c* haplotype) background (named KN-*NAMa*-OE), and *TaNAM-A1d* overexpression lines in wheat variety Fielder (*TaNAM-A1a* haplotype) background (named FD-*NAMd*-OE) respectively. We also cloned full-length *TaNAM-A1d* of 178A comprising 2085 bp native promoter and 1557 bp coding region and delivered into Fielder background (named FD-*NAMd*-COM). In leaves at seedling stage, although *TaNAM-A1* expression in transgenic lines was highly upregulated than in controls lines (Figure S4a), no difference in senescence phenotype was shown between transgenic lines and control lines (Figure S3b-d). In leaves at grain filling stage, dramatically increased expression of *TaNAM-A1* in flag leaves of both KN-*NAMa*-OE and FD-*NAMd*-OE transgenic lines led to significant early-senescence than control lines (Figure 2a-f). While both of the independent FD-*NAMd*-COM transgenic lines showed delayed senescence for 1-2 days with insignificantly higher expression level of *TaNAM-A1* than Fielder (Figure 2d-f). Furthermore, we identified two missense mutations of *TaNAM-A1* in K1107 and K481 from the *Triticum turgidum* cv. Kronos (with *TaNAM-A1c*) TILLING population. These two missense mutations were predicted to have a deleterious effect on the protein function. K1107 contains a splice junction acceptor mutation in the second intron in *TaNAM-A1*, which leads to a predicted 11-residue deletion (Figure S4a), and showed a slight delay in leaf senescence for 1-2 days (Figure 2h). Another TILLING mutant, K481, contains a stop-gained variation at the beginning of the third exon resulting in a premature stop of translation (Figure S4a) and showed a significant delay in leaf senescence for 2-4 days (Figure 2g and 2h). Both mutations did not significantly alter the *TaNAM-A1* expression at mRNA level (Figure 2i). Taking all results together, *TaNAM-A1* is the causal gene for *qGLD-6A*, and functions as a positive regulator of leaf senescence only at reproductive stage. We hypothesized that both *TaNAM-A1a* and *TaNAM-A1d* are functional alleles because their excessive expression accelerates leaf senescence. But *TaNAM-A1d* may be a relatively attenuated allele compared with *TaNAM-A1a*. When mildly expressed in leaves driven by their native promoters, *TaNAM-A1d* slightly delayed the leaf senescence for 1-2 days under the *TaNAM-A1a* background.

**Figure 2.**
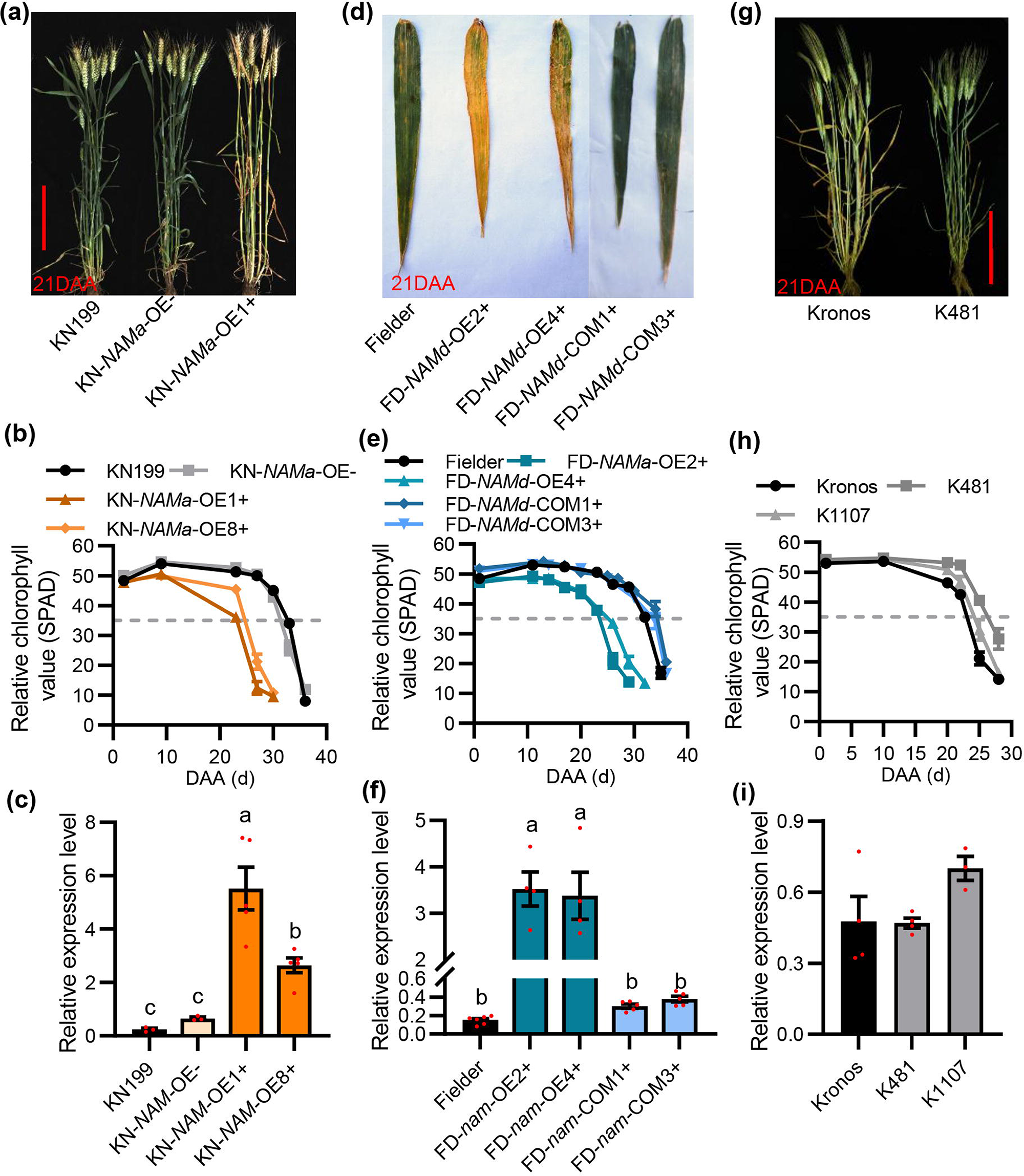
Validation of the *TaNAM-A1* as the major gene of *qGLD-6A* using transgenic lines and TILLING mutants. **(a, d and g)** Comparison of *TaNAM-A1a* **(a)** and *TaNAM-A1d* **(d)** transgenic lines and TILLING mutant K481 **(g)** with their respective control on 21 DAA. Bar = 20 cm. **(b, e and h)** Dynamic SPAD value of flag leaves of *TaNAM-A1a* **(b)** and *TaNAM-A1d* **(e)** transgenic lines and TILLING mutant **(h)** during grain filling stage. Values are the mean ± SE, *n* ≥ 6. Grey dashed line represents the SPAD value = 35. **(c, f and i)** qRT-PCR analysis of *TaNAM-A1* expression level in flag leaves of transgenic lines and TILLING mutants. Values are shown as the mean ± SE, *n* ≥ 3. Different letters indicate significant differences (*P* < 0.05) from a Duncan’s multiple range test.

Interestingly, the increased expression of *TaNAM-A1* in young spikes of transgenic lines (Figure 3a) altered spike length and grain size. The spike length of KN-*NAMa*-OE transgenic lines and FD-*NAMd*-OE transgenic lines were about 25% and 17% longer than that of control (Figure 3b, 3c and 3e). The grains of both KN-*NAMa*-OE and FD-*NAMd*-OE transgenic lines exhibited a remarkable increase in grain length but a strong reduction in grain width and 1000-grain weight (TGW, Figure 3d, 3g-i). FD-*NAMd*-OE transgenic lines produced significantly more grains per spike but no similar effect was observed in KN-*NAMa*-OE transgenic lines (Figure 3f). While the mildly increased expression of *TaNAM-A1d* in FD-*NAMd*-COM imposed less influence on spike length and grain size but produced slightly more grains per spike (Figure 3c, 3d-i). As a result, FD-*NAMd*-COM produced a higher grain yield per spike (GYS) by about 20% than the wild type (Figure 3j) and may possess high yield potential than other transgenic lines. The regulation of TaNAM-A1 on grain size was also reflected among TILLING mutants. K481 and K1107 showed significantly shorter grain length but no significant change in grain width than that of the wild type (Figure S4b-d), and thus the grain weight was slightly decreased in K481 and K1107 than the wild type (Figure S4e). As a summary of the results above, TaNAM-A1 also functions in regulating spike length and grain size. And so far, these findings haven’t been reported before.

**Figure 3.**
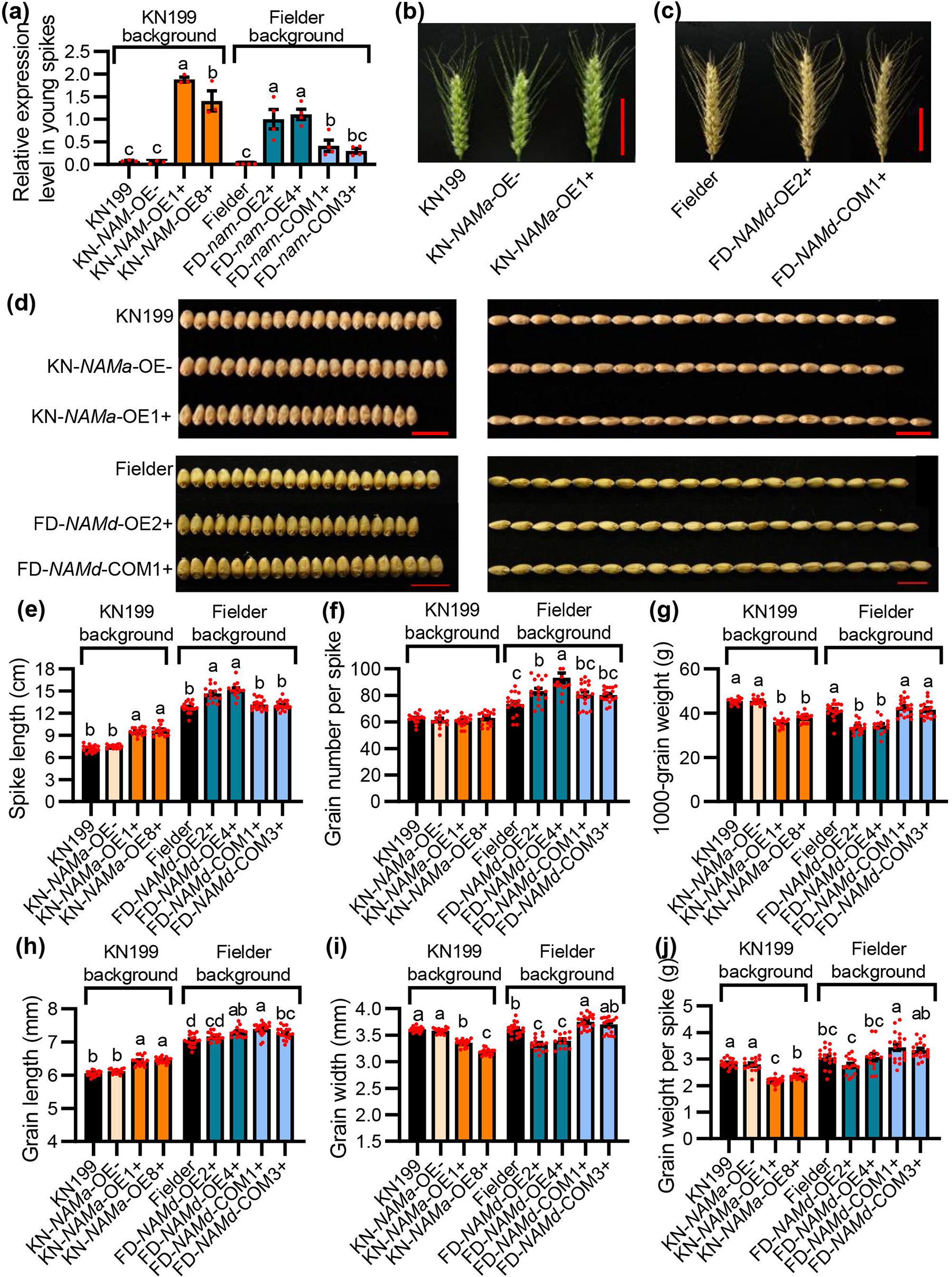
Spike-related and grain-related traits of *TaNAM-A1* transgenic OE lines under field conditions. **(a)** qRT-PCR of *TaNAM-A1* expression level in *TaNAM-A1a* and *TaNAM-A1d* transgenic lines in young spikes at booting stage. Values are shown as the mean ± SE, *n* ≥ 3. Different letters indicate significant differences (*P* < 0.05) from a Duncan’s multiple range test. **(b-c)** Comparison of spike of *TaNAM-A1a* **(b)** and *TaNAM-A1d* **(c)** transgenic lines with control. Bar = 5 cm. **(d)** Comparison of grain size of *TaNAM-A1a* (upper) and *TaNAM-A1d* (down) transgenic lines with control. Bar = 1 cm. **(e-j)** Spike-related traits and grain-related traits of transgenic lines under field condition. Values are shown as mean ± SE, *n* ≥ 14. Different letters indicate significant differences (*P* < 0.05) from a Duncan’s multiple range test.

### Allelic variations of *TaNAM-A1* in 210 Chinese wheat accessions

Based on ecological conditions, growth habbit, and planting time, wheat production in China is divided into ten major agro-ecological production zones (He et al., 2001; Li et al., 2019) (Figure 4f). The Northern Winter Wheat Region (Zone I) and Yellow and Huai River Valleys Facultative Wheat Region (Zone II) are the main growing area for winter wheat and produce 60 to 70% of the total wheat production in China (Qin et al., 2015). We collected 210 modern Chinese wheat varieties from seven provinces in Zone I and II. Three *TaNAM-A1* haplotypes classified by the 731 C to T SNP and the 1518A/deletion polymorphism were found in these 210 Chinese wheat varieties (Figure 4a; Table S4). From north to south in Zones I and II, sowing date of winter wheat changes from late-September to mid-October, while harvest date changes from mid-June to late-May in the next year. As such, the winter wheat crops in the northern provinces have a longer growing season than those in southern provinces. The geographic distribution of the three haplotypes revealed preferences for *TaNAM-A1* by breeders from different areas. In the northern provinces of China, such as Beijing, Hebei, and Shanxi, *TaNAM-A1d* (same as 178A) was the most frequent haplotype among the varieties from these provinces. While in relatively southern provinces such as Shaanxi, Shandong, Henan, and the northern part of Jiangsu and Anhui, *TaNAM-A1a* (same as 178B, China Spring and Fielder) becomes the overwhelming haplotype among the varieties from these provinces (Figure 4g).

**Figure 4.**
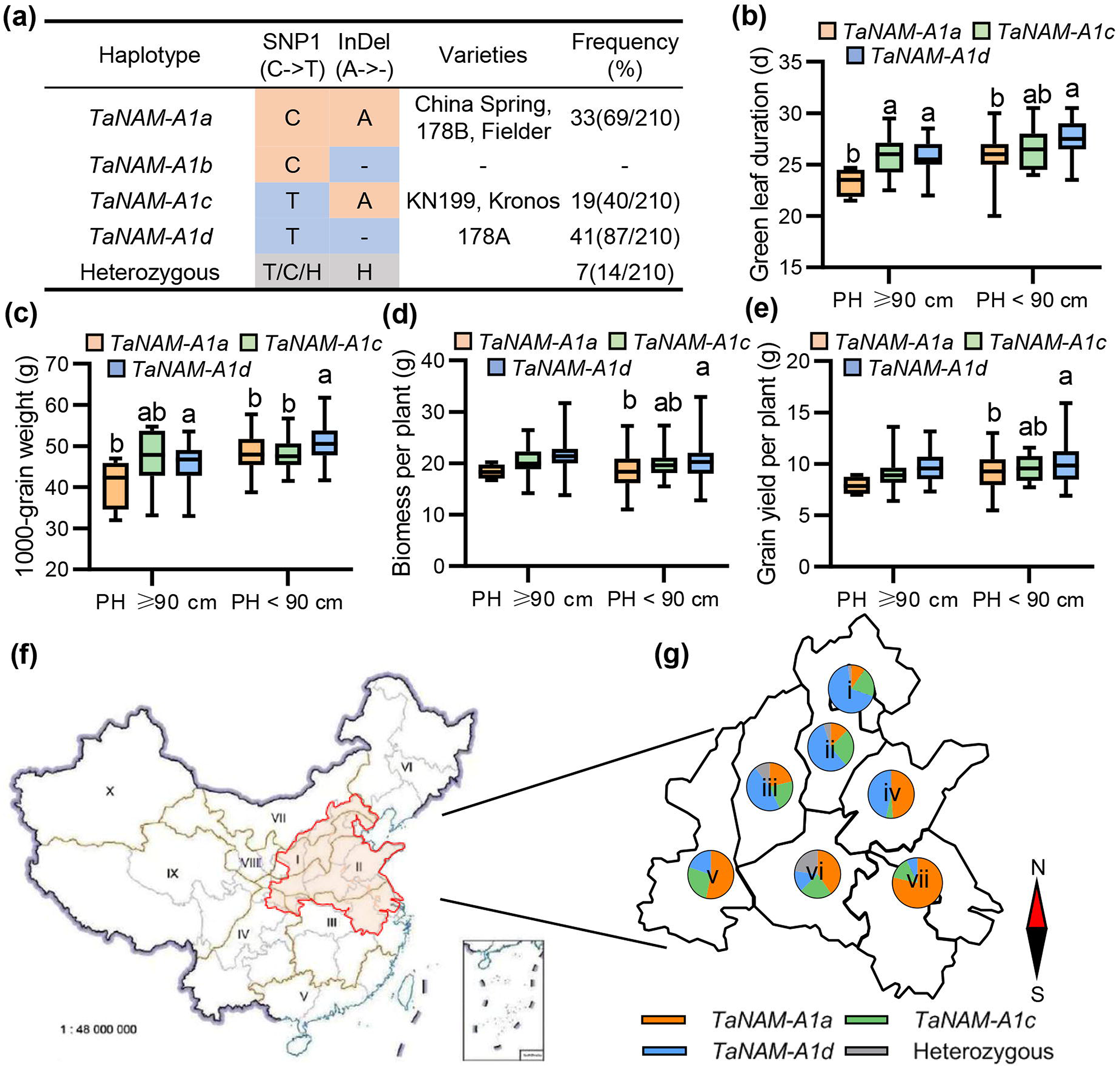
Associations of *TaNAM-A1* haplotypes with green leaf duration and agronomic traits in Chinese wheat varieties. **(a)** Major *TaNAM-A1* haplotypes in 210 Chinese wheat varieties. **(b-e)** Green leaf duration and agronomic traits of three *TaNAM-A1* haplotypes varieties under low nitrogen conditions. Lines in the box plots indicate the median. The 10th/90th percentiles of outliers are shown. Different letters indicate significant differences (*P* < 0.05) from Duncan’s multiple range test. **(f)** Ten agro-ecological zones of wheat in China. I, Northern Winter Wheat Zone; II, Yellow and Huai River Valleys Facultative Wheat Zone; III, Middle and Low Yangtze River Valley Autumn-Sown Spring Wheat Zone; IV, Southern Autumn-Sown Spring Wheat Zone; V, Southwestern Autumn-Sown Spring Wheat Zone; VI, Northeastern Spring Wheat Zone; VII, Northern Spring Wheat Zone; VIII, Northwestern Spring Wheat Zone; IX, Qinghai-Tibetan Plateau Spring-Winter Wheat Zone; X, Xinjiang Winter-Spring Wheat Zone. The map of China is available at http://bzdt.ch.mnr.gov.cn/index/html and the wheat zones are redrawn from Li et al., 2019. **(g)** Geographic distributions of *TaNAM-A1* allelic variations in different provinces of winter wheat production area of China. i, Beijing; ii, Hebei; iii, Shanxi; iv, Shandong; v, Shaanxi; vi, Henan; vii, Jiangsu.

We investigated the GLD and the agronomic traits of 197 varieties under low N and high N conditions. Principal coordinate analysis (PCoA) depicted the GLD of 197 varieties were clustered into separated groups classified by plant height (PH) and *TaNAM-A1* genotypes (Figure S5a), indicating the plant height and *TaNAM-A1* genotypes are closely related to the GLD. The GLD of taller varieties (PH ≥ 90 cm) is shorter than that of semi-dwarf varieties (Figure 4b; Figure S5b); while within the same plant height group, *TaNAM-A1d* haplotype associated with relatively longer GLD, higher TGW, grain yield and shoot biomass than *TaNAM-A1a* haplotype (Figure 4b-e; Figure S5c-e). *TaNAM-A1c* haplotype varieties were intermediate between those *TaNAM-A1a* and *TaNAM-A1d* haplotype varieties. These findings supported the hypothesis supposed by Cormier et al., 2015 that *TaNAM-A1a* may be the strongest functional variant of the *TaNAM-A1* gene by accelerating senescence but decreasing the grain yield, and mainly selected in areas with a shorter growing season while *TaNAM-A1d* may be a weak allele that mainly selected in areas with a longer growing season.

### *TaNAM-A1* regulates the expression of genes involved in biomacromolecules degradation and Fe/Zn remobilization during leaf senescence

To reveal the regulatory mechanism of TaNAM-A1 on leaf senescence, we conducted the transcriptome analysis of leaves of KN-*NAMa*-OE1 positive lines and negative control at 12 days after anthesis (DAA) and 21 DAA. Among 2480 differentially expressed genes (DEGs) between transcriptomes of KN-*NAMa*-OE1 positive lines and negative control at 12 DAA (namely 12DAA-OE vs. 12DAA-WT) and 12824 DEGs at 21 DAA (namely 21DAA-OE vs. 21DAA-WT), the 1315 overlapping DEGs were determined as the key DEGs (Figure S6a; Table S5). 696 up-regulated genes of the 1315 key DEGs exhibited significant enrichment in terms of biomacromolecule metabolic pathways, while 593 downregulated genes were enriched in terms of photosynthesis-related pathways (Figure S6b-c). We found many well-known SAGs that are involved in the chlorophyll degradation pathway (e.g. *TaPAO-4B*, a homolog of *Arabidopsis* gene *PAO*) (Hörtensteiner, 2006) and nucleic acid degradation pathway (e.g. *TaBFN1-2A*, a homolog of *Arabidopsis* gene *BFN1*) (Farage-Barhom et al., 2011), and genes involved in Zn and Fe remobilization (e.g. *TaHMA2like-7A* and *TaIDS3-7A*) (Masuda et al., 2008; Yamaji et al., 2013; Pearce et al., 2014) and amino acid transport (e.g. *TaATL13-2B*, a homolog of rice gene *OsATL13*) (Zhao et al., 2012), were highly upregulated in *TaNAM-A1a* OE lines (Figure S6d; Table S5). On the contrary, *TaNAC-S-7A*, a negative regulator of leaf senescence (Zhao et al., 2015), as well as the genes involved in chlorophyll biosynthesis pathway, were significantly downregulated in *TaNAM-A1a* OE lines. Several genes involved in seed development and plant hormones regulation mechanisms were also differentially expressed between *TaNAM-A1a* OE lines and control (Figure S6d; Table S5). qRT-PCR analysis further confirmed the expression of *TaPAO-4B*, *TaBFN1-2A*, *TaHMA2like-7A*, *TaIDS3-7A*, and *TaATL13-2B* was strongly upregulated in *TaNAM-A1a* OE lines, but strongly downregulated in TILLING mutants (Figure 5a-e). One exception is that the expression of *TaIDS3-7A* was not detected in Kronos and TILLING mutants (Figure 5e). While *TaNAC-S-7A* was dramatically down-regulated in *TaNAM-A1a* OE lines while highly up-regulated in TILLING mutants (Figure 5f). Several binding motifs of ANAC025 transcription factor, the orthologous of TaNAM-A1 in Arabidopsis (Figure S9), were predicted in the 2-kb upstream promoter regions of these genes using JASPAR database (https://jaspar.genereg.net/) (Figure S7), indicating that these genes are potential targets of TaNAM-A1.

**Figure 5.**
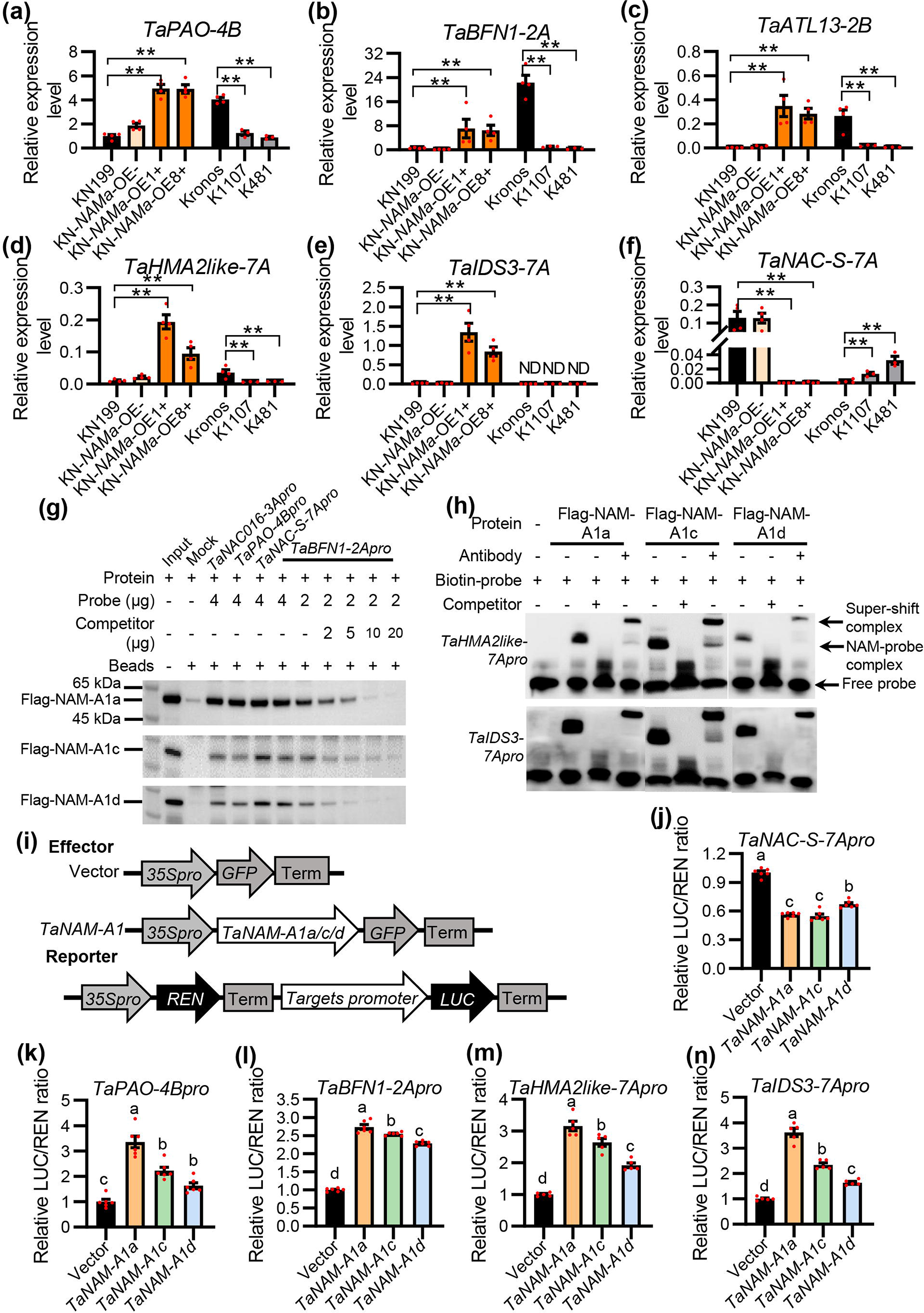
Proteins of three TaNAM-A1 haplotypes show different abilities to transactivate target promoters. **(a-f)** Relative expression of SAGs and genes related to nutrients remobilization in TaNAM-A1 transgenic lines and TILLING mutants by qRT-PCR. Data were represented as mean ± SE, *n* ≥ 3. ** indicates significant difference at *P* < 0.01 compared with the corresponding wild types. **(g)** DNA pull-down assay and **(h)** EMSA showed three TaNAM-A1 proteins directly bind to the promoters of downstream target genes. “+” and “-” indicate the presence and absence respectively. In **(h)**, the amount of competitor probe (unlabeled probe) was 500-fold the amount of biotin-labeled probe. Anti-Flag antibody was used for detecting super-shift complex. **(i)** Schematic diagrams of the effector and reporter plasmids for dual-luciferase transcriptional activity. **(j-n)** Dual luciferase transcriptional activity assay to comparison of the ability of three TaNAM-A1 proteins in transactivating target genes expression. Data were normalized to vector control and represented as mean ± SE, *n* ≥ 5. Different letters denote significant differences (*P* < 0.05) from Duncan’s multiple range test.

### Variations of *TaNAM-A1* haplotypes display diverse transactivation abilities of their encoding protein

Considering natural variations of *TaNAM-1A* contributed to genotypic difference in GLD, overexpression of *TaNAM-A1a* and *TaNAM-A1d* both led to early senescence, we wondered whether there is functional divergence between TaNAM-A1a, TaNAM-A1c and TaNAM-A1d. We firstly surveyed the binding activities of these three proteins to promoter regions of the target genes. DNA pull-down assay and electrophoretic mobility shift assay (EMSA) validated that all of them could directly bind to the promoters of *TaPAO-4B*, *TaBFN1-2A*, *TaHMA2like-7A*, *TaIDS3-7A*, and *TaNAC-S-7A* (Figure 5g and 5h), indicating that the sequence variations of *TaNAM-A1* haplotypes do not affect its recognition and binding activity to core motifs of targets. Then we performed a dual-luciferase transcriptional activity assay to compare their ability to activate target genes expression. All of three proteins significantly activated the expression of *TaPAO-4B*, *TaBFN1-2A*, *TaHMA2like-7A,* and *TaIDS3-7A*. TaNAM-A1a presented the highest transactivation activity while TaNAM-A1d the lowest among the three proteins (Figure 5k-n). On the contrary, all of three proteins significantly repressed *TaNAC-S-7A* expression. The strongest inhibition to *TaNAC-S-7A* expression came from TaNAM-A1a, but the weakest from TaNAM-A1d (Figure 5j). All these results revealed that the sequence variations of *TaNAM-A1* lead to functional divergences in regulating downstream target genes. Also, our results produced sufficient support for the hypothesis that *TaNAM-A1a* is the most functional allele that strongly accelerates senescence by promoting macromolecule degradation in source organs and micronutrient remobilization; while *TaNAM-A1d* has the weakest function in regulating senescence and maintains a longer green canopy duration than other haplotypes.

### TaNAC016-3A, a senescence-induced NAC transcription factor, is a downstream target of TaNAM-A1 and physically interacts with TaNAM-A1 protein

Previous studies have reported that NAC transcription factors often interact with other proteins to form homodimers or heterodimers when binding to target DNA sequences (Olsen et al., 2005; Jeong et al., 2009; Takasaki et al., 2010; Olsen et al., 2015). We carried out a yeast two-hybrid screen to identify the interactive protein of TaNAM-A1. A NAC transcription factor encoded by gene *TraesCS3A02G078400*, was screened as a putative interactive protein with TaNAM-A1 (Figure S8a). TraesCS3A02G078400 is one of the wheat orthologous of rice ONAC016 (LOC_Os01g01430) (Figure S9), so we hereafter named this NAC protein TaNAC016-3A. The following GST pull-down assay, bimolecular fluorescence complementation (BiFC) assay and luciferase complementation imaging (LCI) assay confirmed all three TaNAM-A1 proteins physically interacted with TaNAC016-3A *in vitro* and *in vivo* (Figure 7a-b; Figure S8b-c), indicating that the two mutations in gene coding sequence do not affect the dimerization of TaNAM-A1 protein with TaNAC016-3A.

TaNAC016 has a very close phylogenetic relationship with TaSNAC11 and TaNAC029 (Figure S9), which have been proven to be involved in the regulation of stress tolerance and leaf senescence (Huang et al., 2015; Zhang et al., 2020). As the flag leaves aging during grain filling stage, the expression of *TaNAM-A1* gradually increased at the beginning, reached the peak level at 18 DAA, and then decreased thereafter. While the expression of *TaNAC016-3A* dramatically raised at the late grain filling stage after 18 DAA (Figure 6c). Interestingly, *TaNAC016-3A* also belongs to the 1315 DEGs (Figure S6d), and was significantly upregulated in KN-*NAMa*-OE lines and downregulated in *TaNAM-A1* TILLING mutants (Figure 6d). DNA pull-down assay verified that TaNAM-A1 directly binds to *TaNAC016-3A* promoter (Figure 5g), indicating that *TaNAC016-3A* is a downstream target of TaNAM-A1 and induced by senescence during grain filling stage.

**Figure 6.**
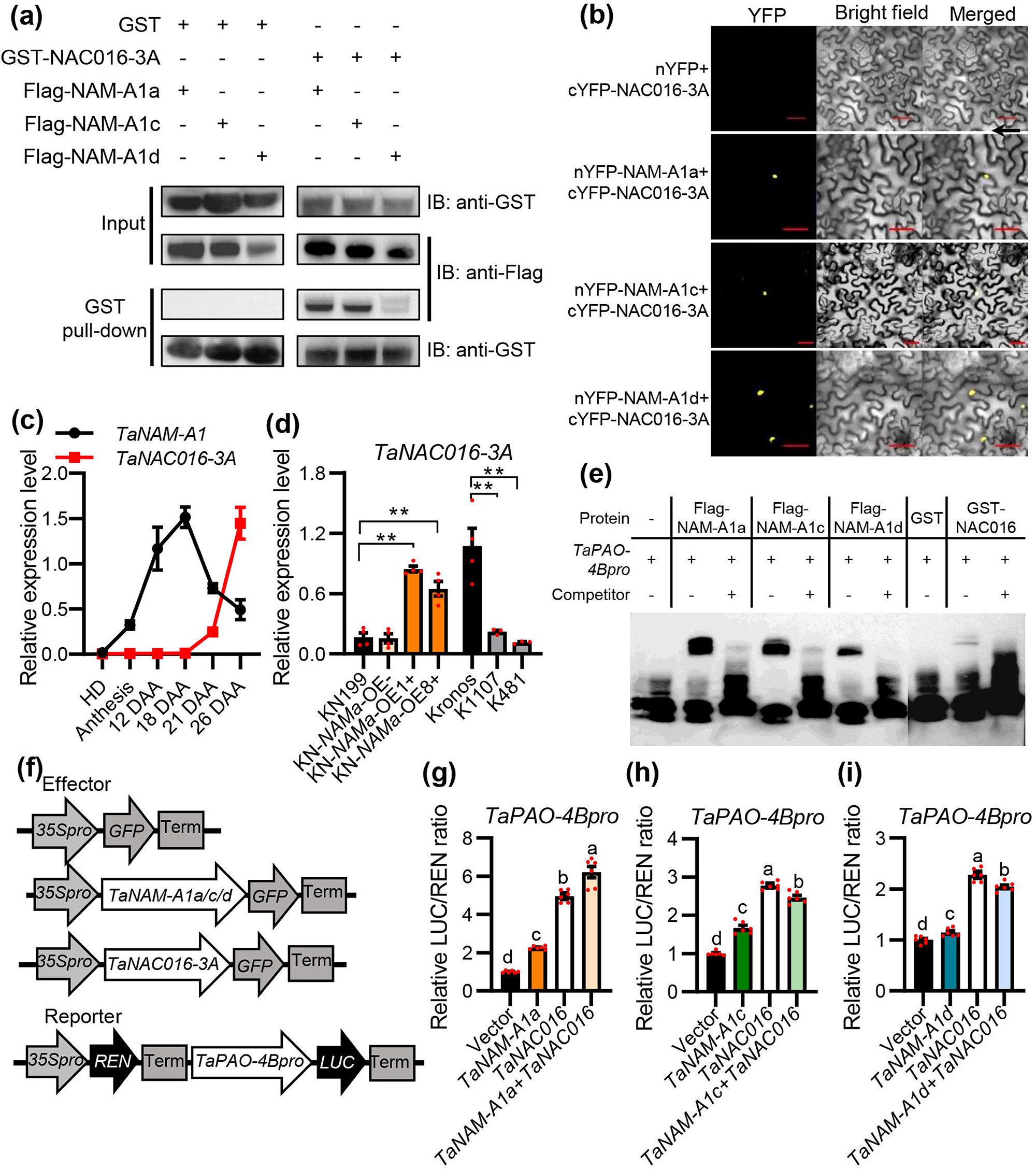
TaNAC016-3A physically interacts with TaNAM-A1. **(a)** GST pull-down assay and **(b)** bimolecular fluorescence complementation (BiFC) assay confirmed three TaNAM-A1 proteins all interact with TaNAC016-3A *in vitro* and *in vivo* respectively. Red scale bar = 20 μm in **(b). (c)** Relative expression of *TaNAM-A1 and TaNAC016-3A* in flag leaves during grain filling stage by qRT-PCR. Values are mean ± SE, *n* = 4. **(d)** Relative expression of *TaNAC016-3A* in transgenic lines and TILLING mutants by qRT-PCR. Data were represented as mean ± SE, *n* ≥ 3. ** indicates significant difference at *P* < 0.01. **(e)** Validation of TaNAM-A1 and TaNAC016-3A directly bind to *TaPAO-4B* promoter by EMSA. “+” and “-” indicate the presence and absence respectively. The amount of competitor probe (unlabeled probe) was 500-fold the amount of biotin-labeled probe. **(f)** Schematic diagrams of the effector and reporter plasmids for dual-luciferase transcriptional activity. **(g-i)** Transcriptional activity assay showed that TaNAC016-3A enhances the transcriptional activation activity of TaNAM-A1 regulating target genes. Data were normalized to vector control and represented as mean ± SE, *n* = 6. Different letters denote significant differences (*P* < 0.05) from Duncan’s multiple range test.

Recent research identified *TraesCS3B02G093300* (named *TaNAC016-3B*), the B genome homeolog of *TaNAC016-3A*, as a candidate senescence regulator and predicted its target genes by GENIE3 network (Harrington et al., 2020). There are several common target genes between the predicted targets of TaNAC016-3B and the key DEGs between *TaNAM-A1a* OE lines and control, including *TaPAO-4B* (Table S5). EMSA validated that TaNAC016-3A, as well as three TaNAM-A1 proteins could directly bind to *TaPAO-4B* promoter (Figure 6e). Subsequently, we performed a dual-luciferase transcriptional activity assay to assess the transactivating activity of the interaction between TaNAM-A1 and TaNAC016-3A to the *TaPAO-4B* promoter. It revealed that the co-expression of TaNAC016-3A with three TaNAM-A1s respectively can highly enhance the transactivating activities to the *TaPAO-4B* promoter than expressing individual *TaNAM-A1* alone (Figure 6f-i). Taking these results together, TaNAC016-3A is also involved in senescence regulation by interacting with TaNAM-A1 and sharing the same target gene with TaNAM-A1. Considering *TaNAC016-3A* is one of the direct downstream targets of TaNAM-A1, and its expression raised only after 18 DAA while the expression of *TaNAM-A1* declined since then, we speculate that TaNAM-A1 triggers senescence initiation and the expression of *TaNAC016-3A*; TaNAC016-3A subsequently takes over the role of promoting senescence in aging leaves.

In addition, we found that co-expression of *TaNAM-A1a* with *TaNAC016-3A* led to significantly higher activity in transactivating *TaPAO-4B* promoter compared with expressing *TaNAC016-3A* alone (Figure 6g). While co-expression of *TaNAM-A1c* and *TaNAM-A1d* with *TaNAC016-3A* led to notably lower transactivating activities than expressing *TaNAC016-3A* alone (Figure 6h-i). Overall, these results indicate that although three TaNAM-A1 proteins all can interact with TaNAC016-3A, their interaction with TaNAC016-3A differentially affects the activity in transactivating downstream genes.

## Discussion

### *TaNAM-A1* haplotypes are functional divergent in regulating leaf senescence during grain filling stage

In this study, we isolated *TaNAM-A1* as the causal gene for the major QTL *qGLD-6A* conferring green leaf duration by map-based cloning. The two mutations at Ala171Val and Gln396fs formed the three TaNAM-A1 proteins in our investigated wheat accessions, TaNAM-A1a (wild type), TaNAM-A1c (with the first mutation only) and TaNAM-A1d (with both mutations). All the three proteins are functional in regulating leaf senescence. Firstly, overexpression of either *TaNAM-A1a* or *TaNAM-A1d* greatly accelerated leaf senescence (Figure 2a-f). TILLING mutations in *TaNAM-A1c* of the *Triticum turgidum* cv. Kronos were shown to delayed leaf senescence in the current study (Figure 2g) and previously (Harrington et al., 2019). Secondly, TaNAM-A1a, -A1c and -A1d all possessed an ability to bind to the promoter regions of *TaPAO-4B*, *TaBFN1-2A*, *TaHMA2like-7A* and *TaIDS3-7A* (Figure 5g and 5h), which are involved in chlorophyll degradation, nucleic acid degradation, Zn and Fe remobilization pathways respectively. However, the three proteins showed discrepant modulating effects on leaf senescence. From a perspective of phenotype, 178B, a parent line used in our map-based cloning, conferred the wild type *TaNAM-A1a* and had about 6 days shorter GLD than the parent line 178A with *TaNAM-A1d* haplotype under both LN and HN conditions (Figure S1b-c). Association analysis on a panel of 197 wheat varieties also revealed that the varieties with *TaNAM-A1d* had significantly longer GLD than those with *TaNAM-A1a* (Figure 4b; Figure S5b). From a perspective of genetic mechanism, the three proteins significantly differed in activities of transactivating senescence-related downstream genes, with TaNAM-A1a the strongest activity and TaNAM-A1d the weakest (Figure 5j-n). Moreover, the protein-protein interaction TaNAM-A1a and TaNAC016-3A enhanced the activity in transactivating *TaPAO-4B*, as compared with TaNAM-A1a or TaNAC016-3A alone (Figure 6g); however, this enhancement was not observed for the interactions between TaNAM-A1c and TaNAC016-3A (Figure 6h) or between TaNAM-A1d and TaNAC016-3A (Figure 6i). As such, *TaNAM-A1d* is a weak allele of *TaNAM-A1* in regulating leaf senescence, and led 178A a longer GLD than 178B. All these results further supported the functional divergences among TaNAM-A1 haplotypes.

### TaNAM-A1 plays a key role in the senescence regulatory pathway involving many senescence-related transcription factors

In this study, we found that the expression of *TaNAM-A1* was hardly detected in the seedling leaves of wild-type wheat, indicating that the regulation of TaNAM-A1 on leaf senescence is age-dependent. The result can be explained by a more recent research which demonstrated TaARF15-A1, an auxin response factor preferentially expressed in young tissues (Table S3), suppressed the expression of *TaNAM-A1* by protein–protein interaction and competition with TaMYC2 for binding to its promoter (Li et al., 2022). In addition, we also found that even though *TaNAM-A1* highly expressed in transgenic lines than in controls lines (Figure S4a), no difference in senescence phenotype was shown between transgenic lines and control lines (Figure S3b-d). This result indicated that there may be unknown factors modulating the function of TaNAM-A1 in senescence initiation before flowering time via multiple-layered regulation including post-transcriptional, translational, and post-translational regulation (Kim et al., 2018).

After flowering stage, the expression of TaNAM-A1 in leaves gradually increased until peaked at 18 DAA, and then sharply decreased in the later days of grain filling stage. This expression pattern is consistent with the findings of Borril et al., 2019, indicating that TaNAM-A1 only expresses and functions at the early stage of grain filling. TaNAM-A1 repressed the expression of *TaNAC-S-7A* (Figure 5f), a negative regulator of senescence and displays decreased expression following leaf senescence (Zhao et al., 2015). We also found that TaNAM-A1 could bind to the promoters of *TaNAC016-3A* and activate its expression at the later stage of grain filling (Figure 5g, 6c-d). Interestingly, Y2H screening followed by BiFC and LCI assays revealed that TaNAM-A1 physically interacted with TaNAC016-3A *in vitro* and *in vivo* (Figure 6a-b; Figure S8a-c). Taking information together, the roles of TaNAM-A1 in regulating leaf senescence are not only triggering biomacromolecules degradation and iron and zinc remobilization, but also being the critical juncture of the senescence regulatory pathway involving many senescence-related transcription factors. Based on the above results, we proposed a working model of TaNAM-A1 in the regulation of leaf senescence (Figure 7a). These findings add new insights into the regulation mechanism of wheat senescence, and further discovering the gene network of senescence-related TFs will promote our understanding.

**Figure 7.**
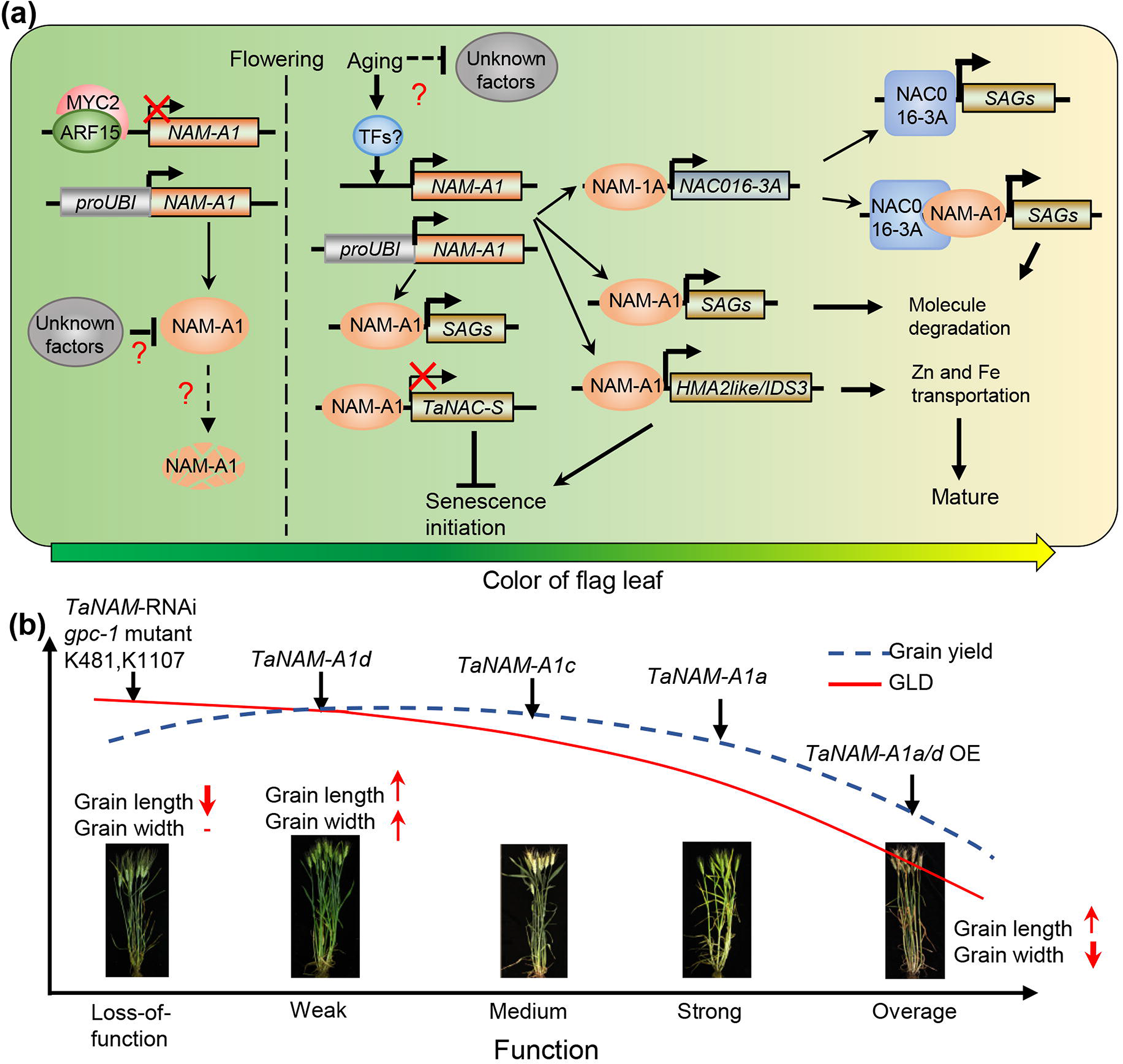
Proposed schematic work models of TaNAM-A1. **(a)** Proposed work model of TaNAM-A1 in regulation of leaf senescence. Before flowering, the expression and function of TaNAM-A1 were repressed by TaARF15-A1 or other unknown mechanism so that it cannot initiate leaf senescence before flowering. After flowering, the repression from the unknown factor is removed. The highly expressed TaNAM-A1 protein binds to downstream target genes promoter, and activates the expression of SAGs and genes involved in micronutrients remobilization, but inhibits the expression of *TaNAC-S-7A*. All these changes trigger the leaf senescence initiation. TaNAM-A1 activates *TaNAC016-3A* expression. Subsequently, TaNAC016-3A interacts with TaNAM-A1, strongly triggers the expression of SAGs, and presumably takes over the role of promoting senescence extensively at the late stage of grain filling. **(b)** A schematic diagram characterizes functional divergences in regulating GLD and yield-related traits during grain filling stage caused by *TaNAM-A1* gene variations and expression level.

### Utilization of *TaNAM-A1* in breeding for balanced stay-green phenotype and yield formation in wheat

Common wheat is the most widely cultivated cereal in the world, providing 20% of the calories and protein in the global human diet. The wheat flag leaf contributes 45-58 % of photosynthetic performance and 41-43 % of assimilates used in grain filling after flowering (Yang et al., 2016). In our study, we found *TaNAM-A1* not only functions as a master regulator of flag leaf senescence, but also involves in regulating spike size and grain size. Increasing *TaNAM-A1* expression had positive effects on spike length, grain number per spike (GNS) and grain length, but had negative effects on grain width and 1000-grain weight, with the degree depending on the *TaNAM-A1* haplotypes overexpressed (the wild type *TaNAM-A1a* vs. the weak allele *TaNAM-A1d*) and the promoters driving *TaNAM-A1* expression (*TaNAM-A1d* driven by ubiquitin promoter vs. by native promoter) (Figure 3). Overexpression of *TaNAM-A1a* and *TaNAM-A1d* both greatly accelerated leaf senescence and significantly reduced TGW (Figure 3g), but overexpressing *TaNAM-A1d* significantly increased GNS (Figure 3f). Consequently, overexpressing *TaNAM-A1a* significantly reduced grain weight per spike (GWS), but overexpressing *TaNAM-A1d* did not (Figure 3j). When *TaNAM-A1d* was driven by its native promoter, it slightly delayed senescence for 1-2 days, had positive effect on GNS (Figure 3f), but did not affect TGW (Figure 3g), and thus increased GWS by 20% compared with the wild type (Figure 3j). Association analysis showed that the wheat varieties harbouring *TaNAM-A1d* had longer GLD and produced about 10% higher value of TGW, biomass and grain yield than those with *TaNAM-A1a* (Figure 4b-e; Figure S5b-e). Association analysis using a panel of European elite varieties revealed *TaNAM-A1d* had an advantage in GNS, TGW and grain yield over *TaNAM-A1a,* and was over-represented in elite varieties (Cormier et al., 2015). As such, *TaNAM-A1d* possesses a longer grain filling duration that is more favorable to yield formation than *TaNAM-A1a* with a shorter grain filling duration.

We proposed a model that characterizes functional divergences in regulating green leaf duration and yield-related traits during grain filling stage caused by *TaNAM-A1* gene variations and expression level (Figure 7b), based on our observed phenotypic differences in a series of genetic materials modifying *TaNAM-A1* function, *TaNAM-A1* haplotypes, and previously reported studies. From the loss-of-function of *TaNAM-A1*, the weak allele *TaNAM-A1d*, the medium allele *TaNAM-A1c*, the wild type *TaNAM-A1a* to overexpression of *TaNAM-A1*, the function of TaNAM-A1 in regulating leaf senescence increases and thus GLD decreases. The presence of *TaNAM-A1a* allele and upregulated expression of *TaNAM-A1* strongly accelerates leaf senescence, resulting in reduced carbon and nutrients accumulation for grain yield formation. The prolonged GLD due to the presence of *TaNAM-A1d* and *TaNAM-A1c* alleles might provide more carbon and nutrients to meet with the assimilates required for a larger spike, resulting in higher grain yield. However, the stay-green phenotype is not always positively correlated with the grain yield formation. The *TaNAM*-RNAi transgenic lines, *gpc-1* mutant and the TILLING mutants reported previously and, in our study, presented an extremely stay-green phenotype but showed no obvious advantages in grain yield to the wild type. It may be explained by the failure of nutrient remobilization from leaves to grains before the end of grain filling stage. This model contributes useful insights for breeding wheat varieties with a balanced extension of leaf functional period and grain yield formation under different geographical climatic conditions.

## Materials and Methods

### Plant materials and field experiments

The 480 recombinant inbred lines (RILs) and two parents 178A and 178B were planted under low N and high N conditions in 2014-2015 and 2015-2016 growing seasons at the experimental station in Beijing and Zhaoxian in Hebei province, respectively. Field identification of the recombinants was conducted in the 2016-2017 growing season under low N conditions at the experimental station in Zhaoxian, Hebei province. Fertilizer application was described previously (Shao et al., 2017). Leaf relative chlorophyll value of each line was measured by SPAD-502 meter (Konica-Minolta, Japan) at different DAA. The green leaf duration (GLD) of each line was used as the phenotype of QTL mapping.

The wild-type KN199 and KN-*NAMa*-OE T2 transgenic lines and their azygous control plants were planted in the field of the experimental station in Zhaoxian, Hebei province in the 2018-2019 growing seasons. The T1 *TaNAM-A1d* transgenic lines and the wild type Fielder were planted in the same location in the 2020-2021 growing season. Young spikes at jointing stage and the flag leaves on 12 DAA and 21 DAA were collected for analysing gene expression at the mRNA level. The SPAD value of the flag leaves at different DAA was investigated to monitor leaf chlorophyll changes. The spike length and grain-size-related traits were characterized after harvest. two *TaNAM-A1* TILLING mutants K481 and K1107 from Kronos TILLING mutant population (Krasileva et al., 2017) were selected to dissect the TaNAM-A1 function. Two dCAPS markers were designed to validate the mutations and the homozygous of the M5 individuals of K481 and K1107 for downstream research. Homozygous mutants of K481 and K1107 growing under high N conditions at the experimental station in Beijing in 2019 and 2020 for phenotyping. Flag leaves on 20 DAA were collected for analysing gene expression at mRNA level.

210 Chinese wheat modern varieties were used for determination of haplotype frequencies and geographic distribution analysis. 197 accessions of the population were planted at the experimental station of China Agricultural University at Quzhou, Hebei province in the 2015-2016 growing season. The GLD and yield-related traits of each accession were characterized. The remaining 13 accessions of Jiangsu province were kindly provided by Prof. Yiwen Li, of which the phenotypes data are unpublished.

### Map-based cloning

DNA samples from 49 lines with extremely long GLD and 49 lines with extremely short GLD, along with 178A and 178B were genotyped by 90K iSelect SNP array (Wang et al., 2014) at Beijing CapitalBio Technology Co., Ltd. 5867 genomic specific and polymorphic SNPs across 21 chromosomes were used for genotyping and genetic map construction (Table S1).The QTL mapping in this study was performed using QTL IciMapping software (Meng et al., 2015). We firstly identified three major regions where abundant GLD-associated SNPs were anchored on chromosomes 1A, 1B and 6A by the Single Marker Analysis (SMA) method, then further dissected the flanking regions of the three underlying QTLs by ICIM-ADD method. The LOD threshold was set to 3.5. For fine mapping of *qGLD-6A*, we developed 9 PCR markers for recombinants screening and genotyping. The primers are listed in Table S7. The publicly accessible gene expression data of wheat developmental tissues were obtained in WheatOmics (http://wheatomics.sdau.edu.cn/) (Pearce et al., 2015;, Ma et al., 2021).

### Vector Construction and Transformation

For wheat genetic transformation, *TaNAM-A1a* CDS with His-tag was cloned into pAHC25 vector (Taylor et al., 1993) to generate *pUbi:TaNAM-A1-His* overexpression construct, which was then transformed into immature embryos of wheat variety KN199 using the method described by Wang et al., 2013. *TaNAM-A1d* CDS with Flag-tag was inserted into pUbiGW vector (Li et al., 2021) to generate the *pUbi:TaNAM-A1d-Flag* overexpression construct. The genomic sequence of *TaNAM-A1d* of 178A comprising 2085 bp native promoter and 1557 bp coding region was also cloned into pUbiGW vector by replacing the *Ubiquitin* promoter of the vector, to generate *pTaNAM-A1d:TaNAM-A1d* complementary construct. Both two constructs of *TaNAM-A1d* were transformed into wheat variety Fielder as described by Li et al., 2019. The primers for vector construction and positive transgenic plant screening were listed in Table S8.

### Transcriptome deep sequencing (RNA-seq)

Total RNA was extracted from flag leaves of KN-*NAMa*-OE1+ and negative control lines at 12 DAA and 21 DAA with three biological replicates using TRIzol reagent (Invitrogen). The RNA-seq libraries construction, transcriptome sequencing and analysis were conducted by Shanghai Oe Biotech Co., Ltd. The RNA-seq reads were aligned to the CS reference genome (International Wheat Genome Sequencing Consortium RefSeq v1.1). FPKM value of each gene and differentially expressed genes (DEGs) was analysed similarly to the method described in (Zhang et al., 2016)

### Quantitative Real-Time PCR

Total RNA treated by DNase I was used for First-strand complementary DNA synthesis with the RevertAid First Strand cDNA Synthesis Kit (Thermo Scientific, K1621). The qRT-PCR analysis was performed with a LightCycler 480 engine (Roche) using Light-Cycler480 SYBR Green I Master Mix (Roche). Relative transcript levels were calculated using the 2^−ΔCT^ method relative to wheat *Ta27922* (Wu et al., 2015). The primers used for qRT-PCR are detailed in Table S8.

### GST pull-down assay

The CDS of *TaNAM-A1a*, *TaNAM-A1c* and *TaNAM-A1d* were cloned into the pETnT vector (Dong et al., 2017) respectively. The full-length CDS of *TaNAC016-3A* were cloned into the pGEX-6p-1 vector. The recombinant Flag-NAM-A1a/c/d and GST-NAC016-3A fusion proteins were expressed in Escherichia coli BL21 (Transseta DE3) with 0.1 mM isopropyl-b-D-thiogalactopyranoside (IPTG) in Luria Bertani (LB) buffer overnight at 16LJ. The GST pull-down assay was performed according to (Gao et al., 2021). Proteins were detected by immunoblot using the anti-Flag and anti-GST antibodies. Each GST pull-down was repeated at least three times. Relevant primer sequences are listed in Table S8.

### DNA pull-down assay

The recombinant Flag-TaNAM-A1a/c/d fusion proteins were expressed in *Escherichia coli* BL21 (Transseta DE3) as described above. Flag-fusion protein purification was performed using Pierce™ Anti-DYKDDDDK Magnetic Agarose (Thermo Scientific, A36797) according to the manufacturer’s instructions.

For DNA–protein pulldown, the probes of target promoters were obtained by PCR amplification with synthetic biotinylated primers. The biotinylated probes were first incubated with Streptavidin Magnetic Beads (MedChemExpress, HY-K0208) in Buffer I (1 M NaCl, 10 mM Tris-HCl, 1 mM EDTA, 0.05% Nonidet P40) for 2 h at 4LJ, then washed three times in Buffer I. DNA-bound beads were then resuspended with Buffer II (100 mM potassium glutamate, 50 mM Tris-HCl pH 7.6, 2 mM MgCl_2_, 0.05% Nonidet P40). Flag-TaNAM-A1a/c/d fusion proteins were then added to DNA-bound beads and the mixture was rotated for 2 h 4LJ. Beads were washed three times with Buffer II. Proteins were stripped off the beads by boiling with 1 × SDS buffer and then subjected to Western Blot. Proteins were detected by immunoblot using the anti-Flag antibody. Each DNA pull-down assay was repeated at least three times. Relevant primer sequences are listed in Table S8.

### Electrophoretic mobility shift assay (EMSA)

The Flag-NAM-A1a/c/d fusion proteins were expressed and purified as described above. The recombinant GST-NAC016 fusion proteins were expressed in Escherichia coli BL21 and purified to homogeneity using GE sepharose 4B beads (GE, Boston, MA, USA) according to the manufacturer’s instructions. EMSAs were performed using the LightShift Chemiluminescent EMSA Kit (Thermo Fisher Scientific, 20148). 200 fmol biotin-labeled probes were added for each reaction. 0.5 μl anti-Flag antibody was added for super-shift complex detection. Each EMSA was repeated at least three times. The sequences of the probes and relevant primers are listed in Table S8.

### Yeast two-hybrid screen

The wheat yeast two-hybrid cDNA library was constructed by Shanghai Oe Biotech Co., Ltd. The full-length and truncate CDS of *TaNAM-A1a* (residues 1–220) were cloned into bait vector pGBKT7 (Clontech) respectively to generate bait plasmids BD-NAM-A1_full_ and BD-NAM-A1_1-220_. BD-NAM-A1_1-220_ with non-autoactivation was used as bait for two-hybrid screening. To validate the interaction between TaNAM-A1 and TaNAC016-3A by one-to-one confirmation experiment, the CDS of *TaNAC016-3A* were fused into the yeast vector pGADT7 (Clontech) to generate AD-NAC016-3A prey plasmid. The yeast two-hybrid and one-to-one confirmation experiments were performed according to Matchmaker™ Gold Yeast Two-Hybrid System User Manual (Clontech, https://www.clontech.com/). Primers for vector construction were listed in Table S8.

### Bimolecular Fluorescent Complimentary (BiFC) assay

The full-length CDS of *TaNAM-A1a*, *TaNAM-A1c* and *TaNAM-A1d* were inserted into the YFN43 (Belda-Palazon et al., 2012) vector, and the CDS of *TaNAC016-3A* was inserted into the YFC43 vector respectively by Invitrogen Gateway® Technology (Thermo Fisher Scientific) to generate nYFP-NAM-A1a/c/d and cYFP-NAC016-3A fusion constructs. Then nYFP-fusion and cYFP-fusion constructs were co-transfected into tobacco leaf epidermal cells by Agrobacterium-mediated infiltration as described previously (Chen et al., 2008). After 48 h incubation in the dark, the YFP signal was examined and photographed using a confocal microscope (Zeiss LSM710). Each BiFC assay was repeated at least three times. Relevant primer sequences are listed in Table S8.

### Luciferase complementation imaging (LCI) assay

LCI assay experiments were conducted according to Chen et al., 2008. The full-length CDS of *TaNAM-A1a*, *TaNAM-A1c*, and *TaNAM-A1d* were inserted downstream of cLUC in 35S::cLUC plasmid, the full-length CDS of *TaNAC016-3A* were cloned to upstream of nLUC in 35S::nLUC plasmid. Derivative NLuc- and CLuc-fusion constructs were co-transfected into tobacco leaf epidermal cells by *Agrobacterium*-mediated infiltration. The LUC images were captured by the low-light cooled CCD imaging apparatus (CHEMIPROHT 1300B/LND, 16 bits; Roper Scientific). Images presented in the figures are representative of six tobacco leaves.

### Dual-luciferase transcriptional activity assay

Dual-luciferase transcriptional activity assays were performed as previously described (Gao et al., 2021). Fragments of 2 kb promoters upstream of *TaPAO-4B*, *TaBFN1-2A*, *TaNAC-S-7A*, *TaHMA2like-7A*, *TaIDS3-7A* were cloned into the pGreenII 0800-LUC vector as reporter plasmids (Hellens et al., 2005). The full-length CDS of *TaNAM-A1a/c/d* and TaNAC016-3A were cloned upstream of the GFP gene in the pCAMBIA1300 vector respectively to generate *p35S::TaNAM-A1a/c/d-GFP* and *p35S::TaNAC016-3A-GFP* as the effector plasmids. *Agrobacterium* (strain GV3101) harboring different combinations of effector and reporter plasmids were co-infiltrated with the same 4-week-old *N. benthamiana* leaves. The Dual-Luciferase Reporter Assay System (Promega, E1960) was used to perform the luciferase activity assay, with the *Renilla* luciferase gene as an internal control. Firefly luciferase activity of the *p35S::TaNAM-A1a/c/d-GFP* and *p35S::TaNAC016-3A-GFP* effectors were normalized to the empty vector control from the same leaves to figure out the relative LUC/REN ratio. At least six biological replicates were used for each dual-luciferase transcriptional activity assay.

### Statistical analysis of data

Data were statistically analysed by two-sided t-test and the multiple comparisons were made using Duncan’s multiple range test by SPSS 22.0 software. *P* values of less than 0.05 were considered to indicate statistical significance. Figures of statistical analysis were drawn using Graphpad Prism 9 software.

## Data Availability

Data supporting the findings of this work are available within the manuscript and its Supplementary Information files. The RNA-seq raw data are available from the National Center for Biotechnology Information Sequence Read Archive (https://www.ncbi.nlm.nih.gov/sra/) with the BioProject accession number PRJNA842904.

## Accession numbers

The gene sequence data from this article can be found in the Chinese Spring reference sequences (IWGSC RefSeq v1.1) under the following accession numbers: *TaNAM-A1* (TraesCS6A02G108300), *TaATL13-2B* (TraesCS2B02G343300), *TaBFN-2A* (TraesCS2A02G507400), *TaPAO-4B* (TraesCS4B02G311100), *TaNAC-S-7A* (TraesCS7A02G305200), *TaNAC016-3A* (TraesCS3A02G078400), *TaHMA2like-7A* (TraesCS7A02G420000) and *TaIDS3-7A* (TraesCS7A02G037700).

## Acknowledgements

We are grateful to Prof. Caixia Gao’s laboratory (Institute of Genetics and Developmental Biology, Chinese Academy of Sciences) for developing the transgenic wheat lines. We also grateful to Prof. Daolin Fu of State Key Laboratory of Crop Biology, Shandong Agricultural University, Tai’an, Shandong China for the gift of the seeds of wild type Kronos and the M5 of K481 and K1107; Prof. Yiwen Li of Institute of Genetics and Developmental Sciences, Chinese Academy of Sciences, Beijing, China for providing the 13 wheat accessions of Jiangsu province. Many thanks to Dr. Genying Li of Shandong Academy of Agriculture Sciences, Jinan, China, Prof. Yongrui Wu of Shanghai Institute of Plant Physiology and Ecology, Chinese Academy of Sciences, Shanghai, China, Prof. Yunhai Li and Dr. Yapei Liu of Institute of Genetics and Developmental Sciences, Chinese Academy of Sciences, Beijing, China for providing vectors for wheat transformation, dual-luciferase transcriptional activity assay, and prokaryotic expression respectively. This research was supported by the Strategic Priority Research Program of Chinese Academy of Sciences (Grant No. XDA24010202).

## Author contributions

Y.T. designed and supervised the project. L.Z. designed and performed most of the fields trials, experiments, data analysis and interpreted the results. C.S., Y.L. and X.Z. contributed to the collection and field trails of wheat materials used in this study. W.T. performed wheat transformation mediated by *Agrobacterium tumefaciens*. C.S. performed data analysis of the Chinese wheat accessions and interpreted the results. W.T., X.H., Y.J. and Z.H. provided technical assistance. L.Z. and Y.T. wrote the manuscript.

## Supplementary information

### Supplementary figures

**Figure S1** QTL mapping for green leaf duration (GLD).

**Figure S2** Sequence variations of *TraesCS02G108100* in 178A and 178B compared with Chinese Spring reference genome.

**Figure S3** Phenotypes of seedlings of *TaNAM-A1a* and T*aNAM-A1d* transgenic lines.

**Figure S4** Grain size of *TaNAM-A1* TILLING mutants compared with the wild type.

**Figure S5** Phenotype analysis of three *TaNAM-A1* haplotypes varieties.

**Figure S6** Transcriptomic analysis of flag leaves of *TaNAM-A1* transgenic lines during grain filling.

**Figure S7** Putative ANAC025 binding motifs in the promoters of *TaPAO-4B*, *TaBFN1-2A*, *TaNAC016-3A*, *TaNAC-S-7A*, *TaHMA2like-7A* and *TaIDS3-7A*.

**Figure S8** TaNAC016-3A physically interacts with TaNAM-A1.

**Figure S9** Phylogenetic tree analysis of TaNAM-A1 and TaNAC016 with other NACs transcription factors showing high similarity in wheat, rice, and Arabidopsis.

### Supplementary tables

**Table S1.** 5867 SNPs used for QTL mapping for GLD.

**Table S2.** Major QTLs for green leaf duration.

**Table S3.** Expression pattern of genes within region of *qGPD-6A* (I∼VII) and *TaARF15*.

**Table S4.** Allelic variations of *TaNAM-A1* in 210 Chinese wheat accessions.

**Table S5.** 1315 DEGs between *TaNAM-A1a* overexpression lines with control lines.

**Table S6.** Protein sequences of NAC transcription factors in wheat, rice and Arabidopsis used for phylogenetic tree analysis.

**Table S7.** PCR markers for marker-assisted selection (MAS).

**Table S8.** Primers used for gene cloning, vectors construction and expression analysis.

